# High-quality complete genome resource of plant pathogenic bacterium *Pectobacterium atrosepticum* strain Green1 isolated from potato (*Solanum tuberosum* L.) in Greenland

**DOI:** 10.1101/2021.06.01.446529

**Authors:** Robert Czajkowski, Lukasz Rabalski, Maciej Kosinski, Eigil de Neergaard, Susanne Harding

## Abstract

*Pectobacterium atrosepticum* is a narrow host range pectinolytic plant pathogenic bacterium causing blackleg of potato (*Solanum tuberosum* L.) worldwide. Till present, several *P. atrosepticum* genomes have been sequenced and characterized in detail; all of these genomes have come, however, from *P. atrosepticum* strains isolates from plants grown in temperate zones, not from hosts cultivated under different climatic conditions. Herewith, we present the first complete, high-quality genome of the *P. atrosepticum* strain Green1 isolated from potato plants grown under subarctic climate in Greenland. The genome of *P. atrosepticum* strain Green1 consists of one chromosome of 4,959,719 bp., with a GC content of 51% and no plasmids. The genome contains 4531 annotated features, including 4179 protein-coding genes (CDSs), 22 rRNA genes, 70 tRNA genes, 8 ncRNA genes, 2 CRISPRs and 126 pseudogenes. We believe that the information of this first, high-quality, complete, closed genome of *P. atrosepticum* strain isolated from host plant grown in subarctic agricultural region will provide resources for comparative genomic studies and for analyses targeting climatic adaptation and ecological fitness mechanisms present in *P. atrosepticum*.

## Genome announcement

*Pectobacterium atrosepticum* (former *Erwinia carotovora* subsp. *atroseptica*), belonging to Soft Rot *Pectobacteriaceae* (SRP; former pectinolytic *Erwinia* spp.), is a plant pathogenic bacterium with a narrow host range primarily limited to potato (*Solanum tuberosum* L.) cultivated in temperate agricultural regions (Harris and Lapwood, 1977; Pérombelon, 2000).

In the past, particularly in Europe and North America, *P. atrosepticum* was regarded as the dominant causative agent of potato blackleg, repeatably isolated from both symptomatic and symptomless potato plants grown in agricultural regions across Europe and worldwide (Perombelon, 1991, 2002). In the 1970s, Molina and Harrison (1977) reported that the high blackleg incidences in potato crop in Colorado, USA, were almost exclusively caused by *P. atrosepticum* (Molina and Harrison, 1977). Perombelon (1972) in Scotland showed that up to 80% of progeny tubers were contaminated with *P. atrosepticum*, although the great majority of infected plants did not show any disease symptoms (Perombelon, 1972). Today, the bacterium still remains the prevalent SRP species causing blackleg in several temperate areas of the Northern Hemisphere, including the U.K., Norway and Canada (van der Wolf et al., 2021). Results from the seed potato surveys performed recently in England, Wales, and Scotland indicated that *P. atrosepticum* still represented the majority (over 89%) of all samples containing pectinolytic bacteria (van der Wolf et al., 2021).

Similarly, *P. atrosepticum* was also one of the most frequently detected *Pectobacterium* species in surveys done between 2015 and 2017 in North Ireland (Zaczek-Moczydłowska et al., 2019). Likewise, some SRP bacteria are also known to live as epiphytes and endophytes on plants and as saprophytes in soil and groundwater (Bell et al., 2004). However, these types of *P. atrosepticum* lifestyle remain largely unexplored yet (Bell et al., 2004). Therefore, it cannot be excluded that the pathogen is more widely distributed in the environment outside agricultural fields in Europe and worldwide than expected.

*P. atrosepticum* is known to be found in cool temperate regions worldwide, causing symptoms at average temperatures below 25 °C (Perombelon, 2002). Temperature plays an essential role in the pathogenicity of SRP bacteria (Lebecka, 2013; du Raan et al., 2015). It may act as a trigger that activates the expression of specific virulence factors during infection (Smirnova et al., 2001). It is widely acknowledged that different SRP species possess different growth temperature optima (Lebecka, 2013; du Raan et al., 2015), under which they can prevail in the environment and cause disease symptoms in host plants. Comparing the optimal growth temperatures of all SRP bacterial species, *P. atrosepticum* isolates are known to grow and cause symptoms in plants at the lowest environmental temperatures (18 °C) observed during crop cultivation (du Raan et al., 2015; Lebecka et al., 2018).

The *P. atrosepticum* strain NCPPB 549 (CFBP 1526, CIP 105192, G/39, ICMP 1526, ICPB EA 172, LMG 2386, PDDCC 1526, genome Genbank accession: JQHK00000000.1) isolated in 1957 in Scotland, UK is recognized as a type strain of this species (Gardan et al., 2003). In contrast, strain SCRI 1043 (ATCC BAA-672, genome Genbank accession: NC_004547.2) isolated in 1985 in the U.K. has been widely used as a reference for the majority of the (molecular) work done to characterize the virulence determinants of the pathogen and *P. atrosepticum* interactions with host plants (Hinton et al., 1989; Mulholland et al., 1993; Toth et al., 1997; Toth et al., 1999a; Bell et al., 2004). To date, several other *P. atrosepticum* genomes have been sequenced and characterized (https://www.ncbi.nlm.nih.gov/genome/1088). However, at the moment of manuscript preparation (May 2021), all of these genome sequences have come from *P. atrosepticum* strains isolated exclusively from plants grown in temperate climate zones.

*P. atrosepticum* strain Green1 was isolated from potato (*Solanum tuberosum* L.) plant in 2019 in Southern Greenland (Neergaard de et al., 2020). During the blackleg and soft rot surveys done in the potato fields in agricultural regions, samples of plants and tubers with typical blackleg and soft rot symptoms were collected at six areas and tested for the presence of SRP bacteria, including *P. atrosepticum*. Ten strains, named Green1 – Green10, representing SRP isolates from symptomatic plants from different locations in Greenland, were all classified as *P. atrosepticum* based on 16S rRNA, *recA*, and detailed biochemical tests (Neergaard de et al., 2020).

It is generally accepted that *P. atrosepticum* strains isolated so far demonstrate a high degree of similarity (de Boer et al., 1979; Toth et al., 1999b; Sledz et al., 2000). Therefore, to assess the genetic diversity of the Green1 – Green 10 isolates collected in the survey, the rep-PCR (Versalovic et al., 1994), executed as described in (Czajkowski et al., 2010), was used. Rep-PCR analysis showed that the fingerprints of all 10 isolates were identical (data not shown).

In our comparative phenotypic studies involving *P. atrosepticum* strains Green1-Green10 and the reference strain SCRI 1043, we detected that Greenlandic strains, but not SCRI 1043, showed the ability to utilize pyruvic acid methyl ester, L-asparagine, myo-inositol and acetic acid. In contrast, SCRI 1043, but not Green1-Green10 isolates, used D-galacturonic acid, D-maltose, D-turanose and D-glucuronic acid as the sole carbon source (Czajkowski, unpublished).

Subarctic and arctic agricultural areas represent a unique environment characterized by extreme temperature, elevated UV radiation, low nutrient and low water content (McCartney and Lefsrud, 2018; Altdorff et al., 2021). This environment may be a source of microbes, including adapted plant pathogens with unique features. The Southern Greenland agricultural regions (60 ° - 61° 15’ N) are characterized by the average temperatures from −7 to - 4 °C in January and from 9 to 11 °C in July and annual precipitations varying between 260 and 815 mm depending on the location (https://www.timeanddate.no/). To further explore the hypothesis that Greenlandic isolates of *P. atrosepticum* may show distinct phenotypic and genetic characteristics from *P. atrosepticum* reference strain SCRI 1043 and to link the preliminarily observed phenotypes with the genomic data, the genome of Green1 strain has been sequenced and annotated.

For the isolation of genomic DNA, *P. atrosepticum* strain Green1 was grown at 28 °C in tryptone soya broth (TSB, Oxoid) for 16 h with shaking (150 rpm). Bacterial DNA was isolated using a Wizard Genomic DNA Purification Kit (Promega) according to the protocol provided by the manufacturer. The Green1 genome was sequenced by a hybrid approach with long reads generated using Oxford Nanopore Technologies (ONT) and short reads generated using Illumina technology, as described earlier (Czajkowski et al., 2021). DNA sequencing libraries were prepared according to the manufacturer (for ONT, it was LSK kit, and for Illumina, it was a Nextera XT kit). Long reads were generated by a single Flongle run on MinION, and short reads were generated during the Midoutput run on Ilumina Miniseq. *De novo* assembly of long reads was performed and validated using Pomoxis (https://nanoporetech.github.io/pomoxis/), Medaka (https://github.com/nanoporetech/medaka) and Unicycler (https://github.com/rrwick/Unicycler) with final mean coverage x 230. Initial genome polishing was conducted by a tool integrated into the assembler. Pilon 1.23 (https://github.com/broadinstitute/pilon) was used to further error corrections. The combined procedure produced a single contig of 4,959,719 bp. which was further manually inspected and curated. Repetitive genetic elements were searched using Geneious (Kearse et al., 2012). Finally, BLAST (http://blast.ncbi.nlm.nih.gov/Blast.cgi), InterProScan (http://www.ebi.ac.uk/Tools/pfa/iprscan/) and HMMER (phmmer, UniProtKB) (Finn et al., 2011) (http://hmmer.org/) were used to annotate the *P. atrosepticum* Green1 genome. The obtained structural and functional annotations were retested using RAST (Rapid Annotation using Subsystem Technology (http://rast.nmpdr.org/) (Aziz et al., 2008) DFAST (http://dfast.ddbj.nig.ac.jp/) (Tanizawa et al., 2018). Preliminary comparative genomics analyses (Green1 vs. SCRI 1043 (reference strain)) were done using EDGAR 3.0 (https://edgar3.computational.bio.uni-giessen.de) (Blom et al., 2009; Blom et al., 2016). The presence of prophages (bacteriophage sequences) in the genome of Green1 was predicted using Prophage-Hunter (https://pro-hunter.genomics.cn/) (Song et al., 2019) and PHASTER (https://phaster.ca/) (Arndt et al., 2016). The presence of gene clusters involved in the biosynthesis of secondary metabolites was predicted using the antiSMASH 5.0 database (https://antismash.secondarymetabolites.org/#!/start) (Blin et al., 2019).

The complete, closed, high-quality genome of *P. atrosepticum* strain Green1 has been uploaded to NCBI Genbank and received the accession number: CP055224. Upon submission to NCBI Genbank, the genome was also annotated by the NCBI Prokaryotic Genome Annotation Pipeline (https://www.ncbi.nlm.nih.gov/genome/annotation_prok/) (Tatusova et al., 2016). The genome (Fig. 1) is a double-stranded 4,959,719 bp. DNA sequence, possessing 4531 annotated features including 4179 protein-coding genes (CDSs) and an average GC content of 51% with no plasmids. The average gene length was predicted to be ca. 1100 bp. and the average protein length was ca. 366 amino acids. 85.3% of the genome consists of coding regions. The *P. atrosepticum* Green1 genome encodes 22 rRNAs, 70 tRNAs, 8 ncRNA genes, 2 CRISPRs and 126 pseudogenes.

**Figure 1.**
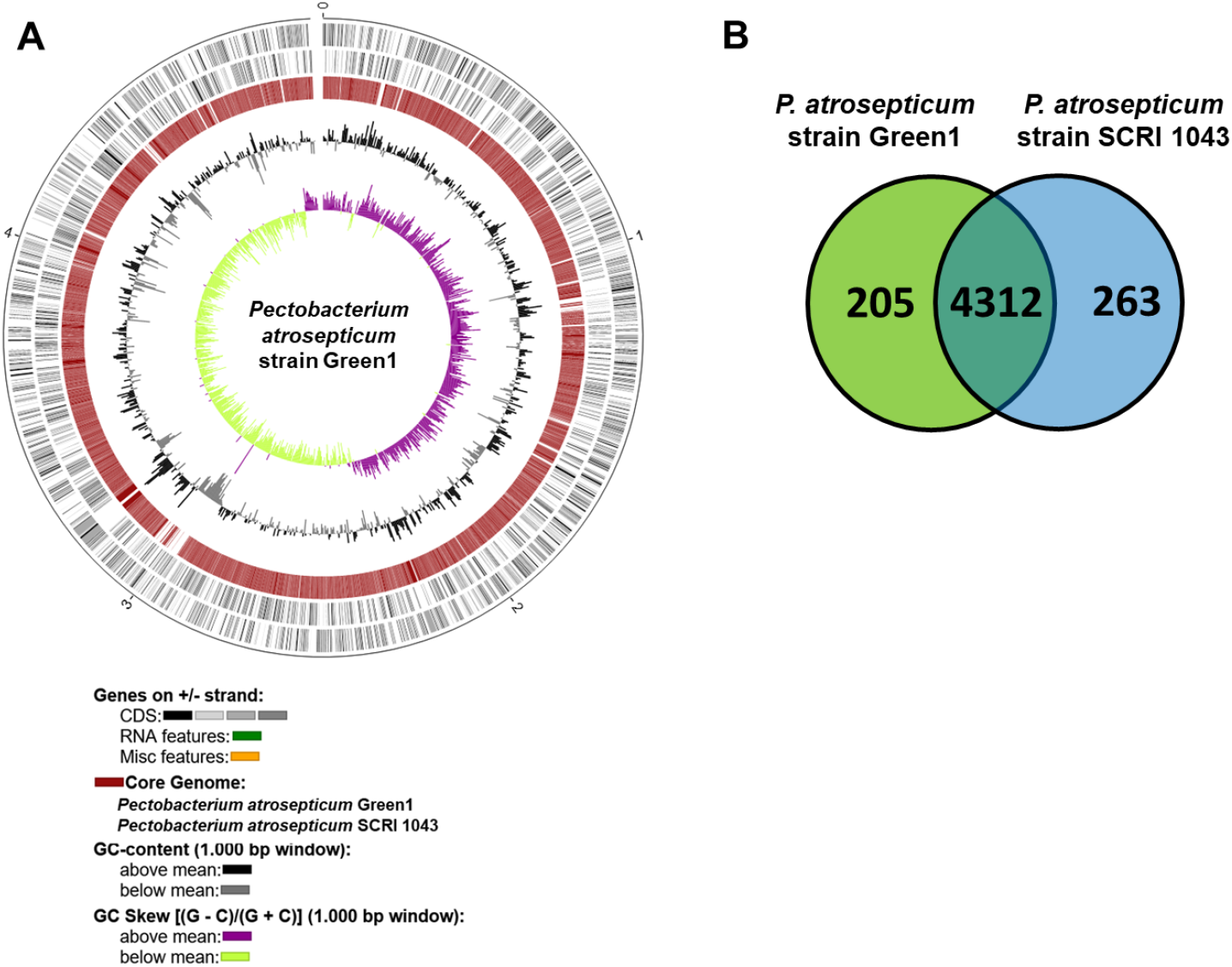
Features of the *P. atrosepticum* strain Green1 genome and its comparison with the SCRI 1043 genome. The circular plot of the Green1 and SCRI 1043 genomes (A) and core genome shared between *P. atrosepticum* strains Green1 and SCRI 1043 (B). The comparative analyses were done using EDGAR 3.0 – an enhanced software platform for comparative gene content analyses (Blom et al., 2009; Blom et al., 2016).

Using antiSMASH, we also identified 9 gene clusters responsible for the synthesis of secondary metabolites in the genome of *P. atrosepticum* strain Green1. They matched the secondary metabolite clusters described previously in *Enerobacteria* (Mohite et al., 2019) and *Pectobacteria* (Jonkheer et al., 2021). Seven of these, *viz*. siderophore, betalactone, phenazine, oronofacic acid, amonabactin, hserlactone and arylpolyene, were present both in the genome of the reference *P. atrosepticum* strain SCRI1043 and Greenlandic strain Green1. Contrary, the other two (anikasin and thiopeptide) were found only in the genome of strain Green1. One of the secondary metabolites predicted to be encoded by the Green1 genome: anikasin, is a cyclic lipopeptide, enabling swarming motility of *Pseudomonas fluorescence* strain HKIO770, protecting the bacterium from protozoal grazing (Gotze et al., 2017). The role of anikasin in the ecology of *Pectobacterium* spp. including *P. atrosepticum* has not been assessed yet, but it can be speculated that the metabolite may protect the Green1 strain from predators found in soil. The second: thiopeptide is an antibiotic known to be produced by microbial dwellers present in soil (Just-Baringo et al., 2014). Likewise, its role in the ecology of *P. atrosepticum* strain Green1 remains unknown. Still, based on literature data, it can be speculated that this antibiotic may be crucial for the competitive survival of the bacterium in the soil environment.

Although the core genome shared between Green1 and SCRI 1043 calculated with EDGAR 3.0 contains 4312 ORFs, the genome of Green1 contains 205 singletons, absent in the genome of *P. atrosepticum* strain SCRI 1043. The great majority of these singletons (168 genes) were predicted to encode hypothetical proteins for which no homology with known proteins could be found. However, the genome of the strain Green1 also contained singletons coding among others for: toxin/antitoxin proteins (CcdA/CcdB), conjugal transfer protein TraA, transcriptional factors, site-specific recombinases, mobile genetic elements and restriction/modification systems not found in the genome of the reference *P. atrosepticum* strain SCRI 1043. These genes may give the bacterium ecological advantages over SCRI 1043 when both species are present in the same niche. Still, more work is needed to (experimentally) assess the (comparative) ecology of *P. atrosepticum* strain Green1 in detail.

The investigation of the Green1 genome for the presence of viral sequences done with Prophage-Hunter and PHASTER did not result in both cases in the prediction of regions containing intact (complete) viral genomes. However, several regions in the genome of Green1 were found, harbouring incomplete prophage genomes or individual phage-derived genes. For example, the gene coding for Tin protein present in the genome of Green1 strain may provide the bacterium with resistance against T-even phages in the environment (Christie and Calendar, 2016), which would have an ecological relevance in the phage-full settings.

The first, completed, high-quality genome of *P. atrosepticum* strain Green1 isolated from potato grown in the subarctic agricultural area may prove helpful in (comparative) climatic adaptation studies and ecological fitness analyses done to understand better the origin, spread and virulence traits of *Pectobacterium* species.

**Table 1.** General features of the *Pectobacterium atrosepticum* strain Green1 complete genome.

|  | **Green1 genome** |
| --- | --- |
| **Genome size (bp.)** | 4,959,719 |
| **G.C. content (%)** | 51% |
| **No. contigs** | 1 |
| **N50** | 1 |
| **Coding ratio (%)** | 85.3% |
| **No. total annotated sequences (features)** | 4531 |
| **Average gene length (bp.)** | 1100 |
| **Presence of plasmids** | no |
| **rRNAs** | 22 |
| **tRNAs** | 70 |
| **CRISPRs** | 2 |
| **No. intact prophage genomes (according to PHASTER and Prophage-Hunter)** | 0 |
| **GenBank accession number** | CP055224 |

## Authors’ statement

The authors declare that no conflict of interest exists.

## Acknowledgements

The work was financially supported by Polish Ministry of Science and Higher Education (Ministerstwo Edukacji i Nauki, Polska) funds DS 531-N104-D800-21 to Robert Czajkowski and by Den Grønlandske Fond - grant No 2018-02-070 / RIGS-GL 2018-542 to Eigil de Neergaard. Non-financial support (transport etc.) was provided by The Agricultural Consulting Services (Greenland).

